# Human gut *Faecalibacterium prausnitzii* deploy a highly efficient conserved system to cross-feed on β-mannan-derived oligosaccharides

**DOI:** 10.1101/2020.12.23.424282

**Authors:** Lars J. Lindstad, Galiana Lo, Shaun Leivers, Zijia Lu, Leszek Michalak, Gabriel V. Pereira, Åsmund K. Røhr, Eric C. Martens, Lauren S. McKee, Sylvia H. Duncan, Bjørge Westereng, Phillip B. Pope, Sabina Leanti La Rosa

## Abstract

β-Mannans are hemicelluloses that are abundant in modern diets as components in seed endosperms and common additives in processed food. Currently, the collective understanding of β-mannan saccharification in the human colon is limited to a few keystone species, which presumably liberate low-molecular-weight mannooligosaccharide fragments that become directly available to the surrounding microbial community. Here we show that a dominant butyrate-producer in the human gut, *Faecalibacterium prausnitzii*, is able to acquire and degrade various β-mannooligosaccharides (β-MOS), which are derived by the primary mannanolytic activity of neighboring gut microbiota. Detailed biochemical analyses of selected protein components from their two β-mannooligosaccharides (β-MOS) utilization loci (*Fp*MULs) supported a concerted model whereby the imported β-MOS are stepwise disassembled intracellularly by highly adapted enzymes. Coculturing experiments of *F. prausnitzii* with the primary degrader *Bacteroides ovatus* on polymeric β-mannan resulted in syntrophic growth and production of butyrate, thus confirming the high efficiency of the *Fp*MULs’ uptake system. Genomic comparison with human *F. prausnitzii* strains and analyses of 2441 public human metagenomes revealed that *Fp*MULs are highly conserved and distributed worldwide. Together, our results provide a significant advance in the knowledge of β-mannans metabolism and the degree to which its degradation is mediated by cross-feeding interactions between prominent beneficial microbes in the human gut.

**Importance:** Commensal butyrate-producing bacteria belonging to the Firmicutes phylum are abundant in the human gut and are crucial for maintaining health. Currently, insight is lacking into how they target otherwise indigestible dietary fibers and into the trophic interactions they establish with other glycan degraders in the competitive gut environment. By combining cultivation, genomic and detailed biochemical analyses this work reveals the mechanism enabling *F. prausnitzii*, as a model clostridial cluster IV Firmicute, to cross-feed and access β-mannan-derived oligosaccharides released in the gut ecosystem by the action of primary degraders. A comprehensive survey of human gut metagenomes shows that *Fp*MULs are ubiquitous in human populations globally, highlighting the importance of microbial metabolism of β-mannans/β-MOS as a common dietary component. Our findings provide a mechanistic understanding of the β-MOS utilization capability by *F. prausnitzii* that may be exploited to select dietary formulations specifically boosting this beneficial symbiont, thus butyrate production, in the gut.

## INTRODUCTION

The human distal gut supports a densely populated microbial community that extends the metabolic capabilities lacking in the hosts genome (1). In particular, recalcitrant glycans that are resistant to human digestive enzymes are broken down by the colonic microbiota to monosaccharides and further fermented into host-absorbable short-chain fatty acids (SCFAs). Microbial-borne SCFAs serve critical functions both as energy source, regulators of inflammation, cell proliferation and apoptosis (2). Therefore, catabolism of complex dietary carbohydrates reaching the distal part of the gastrointestinal tract has a central role in shaping the structure and metabolic output of the human gut microbiota and, in turn, host health status (3).

Members of the Gram-positive Firmicutes and the Gram-negative Bacteroidetes phyla constitute the majority of the bacteria found in this ecosystem (4), individual species of which have evolved different strategies to harvest energy from the available dietary glycans (5). Within the Bacteroidetes, *Bacteroides* spp. have been extensively investigated with respect to carbohydrate degradation and they are considered generalists, displaying broad plasticity for glycan utilization (5). *Bacteroides* spp. are particularly notable for dedicating large proportions of their genome to carbohydrate utilization, organizing genes coding for functionally related carbohydrate active enzymes (CAZymes), transport and regulatory proteins into polysaccharide utilization loci (PULs) (6). Despite variations in the polysaccharide they target, the key feature of a PUL is the presence of one or more TonB-dependent receptor (SusC-homolog) and a contiguous substrate binding lipoprotein (SusD-homolog). Compared with *Bacteroides*, Firmicutes encode a lower proportional number of CAZymes and are thought to be nutritionally specialized for selected glycans (1, 5). Recently, species within the Firmicutes phylum have been shown to organize cohorts of genes encoding glycan-utilization systems into loci and being primary degraders of common dietary carbohydrates (7-9). Firmicutes typically utilize glycan-specific ATP-binding cassette (ABC) transporters, which mediate high-affinity capture of oligosaccharides via their extracellular solute-binding proteins (SBPs) (5).

*Faecalibacterium prausnitzii*, a member of the clostridial cluster IV within the Firmicutes phylum, is one of the three most abundant species detected in the human gut microbiota and one of the main sources of butyrate in the colon (10). A growing body of evidence recognizes the crucial role played by *F. prausnitzii* populations in maintaining local and systemic host-health as they are often found to be less abundant in individuals affected by colorectal cancer (11) and certain forms of inflammatory disorders, including alternating-type irritable bowel syndrome (IBS), irritable bowel diseases, celiac disease, obesity and type 2 diabetes, appendicitis and chronic diarrhea (12, 13). In addition, studies in mice models have demonstrated that both cell and supernatant fractions of *F. prausnitzii* reduces the severity of acute, chronic and low-level chemically-induced inflammations (14) (15). *F. prausnitzii* also contributes to colonic epithelial homeostasis by stimulating the production of mucin *O*-glycans and by maintaining appropriate proportions of different cell types of the secretory lineage (16). Collectively, these aforementioned properties make *F. prausnitzii* a potential novel health-promoting probiotic (17) and interventions aimed at increasing the representation of these butyrate-producing bacteria may be used to confer protection against several intestinal disorders.

A common component of the human diet are β-mannans. These complex plant glycans are found in high concentrations as naturally occurring dietary fibers in certain nuts, beans, legume seeds, tomato seeds, coconut and coffee beans (18). In addition, mannan hydrocolloids, including guar gum and carob galactomannan (CGM) as well as konjac glucomannan (KGM), are widely used in the food industry to improve the rheological properties of processed products (19). The constant exposure of the gut bacterial community to dietary mannans is consistent with the finding that β-mannan metabolism is one of the core pathways in the human gut microbiota (20). Structurally, β-mannans display source-related diversity with respect to presence of β-1,4-linked mannosyl and glycosyl residues, α-1,6-linked galactosyl groups and acetyl decorations at positions *O*-2, *O*-3, and/or *O*-6 (18). PULs degrading homopolymeric mannan and galactomannan have been described in the glycan generalists *Bacteroides fragilis* and *Bacteroides ovatus*, respectively (21, 22). We recently reported the characterization of a novel β-mannan utilization locus conferring *Roseburia intestinalis*, a model for the clostridial cluster XIVa Firmicutes, with the ability to ferment this fiber through to butyrate via a selfish mechanism (7). β-mannan degradation was proven to be initiated by an endo-acting multi-modular GH26 enzyme localized on the cell surface; the resulting oligosaccharides are imported intracellularly through a highly-specific ABC-transporter, and completely de-polymerized to their component monosaccharides by an enzymatic cocktail containing carbohydrate esterases, β-glucosidases and phosphorylases (7).

Although *F. prausnitzii* has been described as an efficient degrader of host-derived and plant glycans (23), the ability of this important butyrate-producing microbe to utilize dietary β-mannans has received little attention. In a previous study, we reported that wood-derived acetylated galactoglucomannan stimulates the proliferation of *F. prausnitzii* populations in a pH-controlled batch culture fermentation system inoculated with healthy adult human feces (24). However, the molecular mechanism underlining β-mannan utilization by *F. prausnitzii* in the human gut has not been explored to date.

In this study, we describe and biochemically characterize components of two loci that mediate acquisition and catabolism of β-mannooligosaccharides (β-MOS) by *F. prausnitzii* SL3/3. Together, these data allowed us to outline a pathway for dietary β-MOS deconstruction and saccharification to monosaccharides through cross-feeding with *Bacteroides* species, which contributes to the ecology of β-mannan utilization in the gut ecosystem. Remarkably, we show that the binding proteins that confers β-MOS capture in *F. prausnitzii* targeted ligands with stronger affinity than that of *Bacteroides* species, thus providing *F. prausnitzii* with the ability to cross-feed on the β-MOS available in the environment with high efficiency.

## RESULTS

Genes encoding enzymatic activities required to catabolize mannans were identified within two putative mannan utilization loci (MULs) in *F. prausnitzii* SL3/3. The large MUL (*Fp*MULL) consists of fourteen genes encoding nine enzymes, the components of an ABC transporter, a predicted LacI-type transcriptional regulator (TR), and a hypothetical protein (Fig. 1a). The enzymes encoded by *Fp*MULL include an α-galactosidase belonging to the glycoside hydrolase (GH) family 36 (*Fp*GH36), two carbohydrate esterases (CEs, *Fp*CE2 and *Fp*CE17), a GH113 (*Fp*GH113), one epimerase (*Fp*Mep), a β-1,4-mannooligosaccharide phosphorylase (*Fp*GH130_2), a mannosylglucose phosphorylase (*Fp*GH130_1), a phosphomutase (*Fp*Pmm) and a GH1 isomerase (*Fp*GH1). In addition, based on the similarity with *Ri*GH3A and *Ri*GH3B from the previously characterized β-mannan utilization system in *R. intestinalis* (7), genes encoding two predicted GH3 β-glucosidases were identified (*Fp*GH3A and *Fp*GH3B). These two genes are located in a different locus in the genome, hereafter referred to as *Fp*MULS, and are likely to be involved in (galacto)glucomannan turnover. Based on known activities within GH families, the β-1,4-mannan backbone is predicted to be hydrolyzed by extracellular GH26, GH5 and/or GH134 enzymes (see www.cazy.org). However, no gene coding for such enzyme was identified in the genome of *F. prausnitzii* SL3/3. In addition, endo-β-1,4-mannanase activity was originally reported for two GH113 (see www.cazy.org) although we demonstrated that a GH113 within the mannan utilization locus of *R. intestinalis* is a reducing end mannose-releasing exo-oligomannosidase. A gene encoding a GH113 was detected in the *Fp*MULL (Fig. 1a). Based on a genomic context analysis and *in silico* prediction of a signal peptide, the function of *Fp*GH113 would be as an intracellular mannanase or mannosidase, thus its enzymatic function could not be assigned before an in-depth biochemical characterization (see later results for *Fp*GH113).

**Figure 1.**
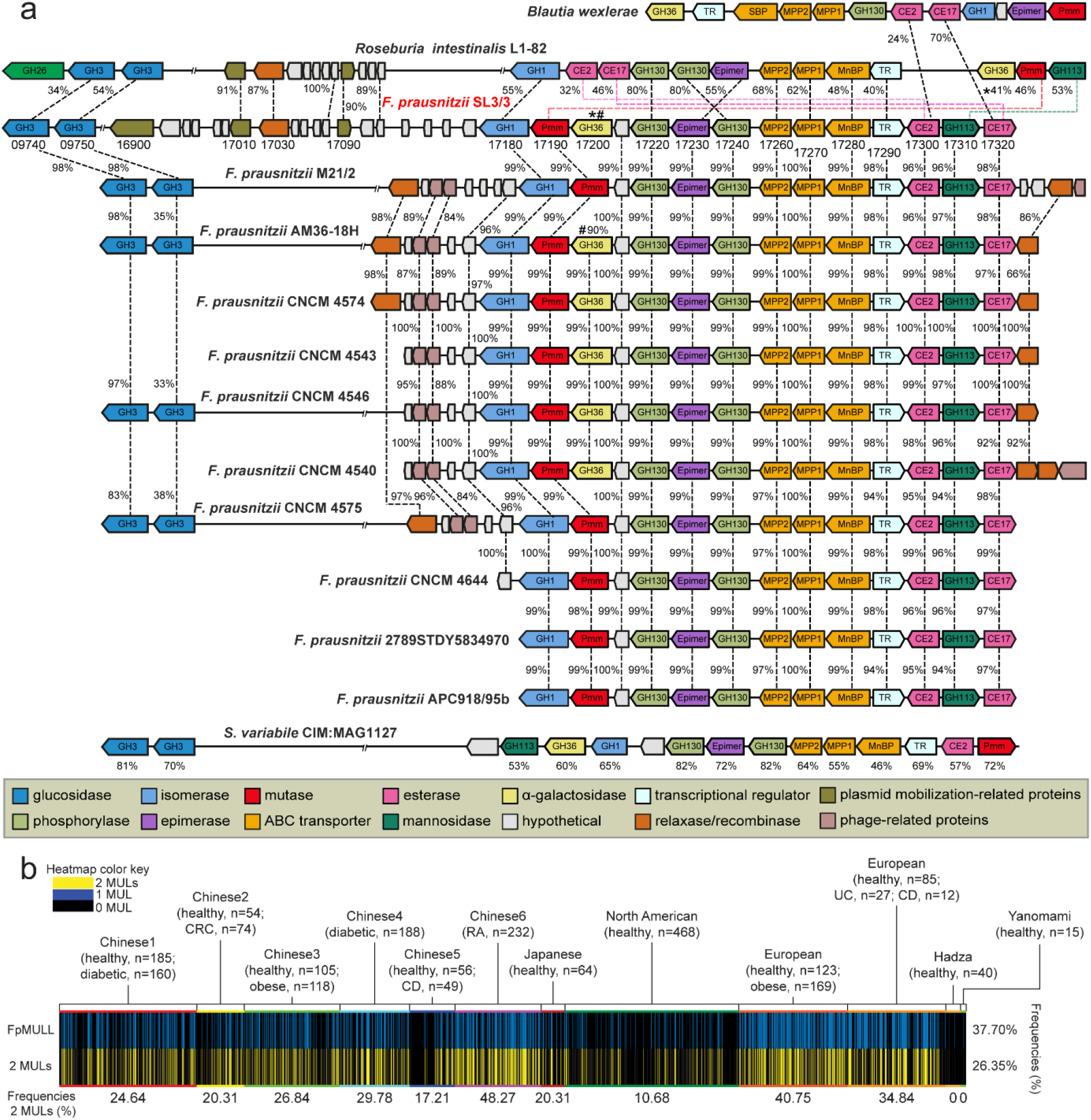
**a**, Large and small β-mannooligosaccharide utilization loci (MULL and MULS, respectively) genomic structure in *F. prausnitzii* SL3/3 and across other publicly available *F. prausnitzii* genomes. In *F. prausnitzii* SL3/3, locus tag numbers FPR_XXXXX are abbreviated with the last numbers after the hyphen. *Fp*MULS corresponds to the genes 09740-09750 while *Fp*MULL includes the genes 17180-17320. Numbers between each gene indicate the percent identity of the encoded protein. Numbers below *S. variabile* CIM:MAG1127 genes indicate the identity of the encoded proteins shared with the same protein in *F. prausnitzii* SL3/3. **b**, Prevalence of *Fp*MULL as well as both *Fp*MULL and *Fp*MULS (2 MULs) in human metagenomes. Each line denotes the presence (blue or yellow) or absence (black) of the *Fp*MULL/MULs related in a single human gut metagenomic sample. The numbers below the bottom row represent the frequency of *Fp*MULs that each cohort possesses. The frequency of *Fp*MULL/MULs incidence across all 2441 individuals is shown on the right.

Genomic comparisons showed that homologous systems to the *Fp*MULL and *Fp*MULS occur in other sequenced *Faecalibacterium* members, with high percentage of similarity (Fig. 1a). Comparison of the gene organization and protein sequence also revealed various levels of rearrangements and moderate protein homology with the two β-mannan utilization loci from *R. intestinalis*. Examination of the regions flanking the *Fp*MULL of *F. prausnitzii* SL3/3 showed the presence of genes encoding plasmid mobilization-related proteins, including a cell invasion protein, a relaxase MobA/VirD2 and a DNA ligase (Fig. 1a). Interestingly, *R. intestinalis* L1-82 genome harbors a similar region including genes coding for the same plasmid-related components, suggesting that the origin of *Fp*MULL could be the result of vertical transfer through bacterial conjugation within colonic microbes. Further comparisons revealed that the genes located upstream and downstream the *Fp*MULL of *F. prausnitzii* M21/2 and six other sequenced *F. prausnitzii* strains code for an incomplete prophage, including one-two relaxases and an integrase (Fig. 1a). These results indicate that phage-related horizontal gene transfer was an alternative mechanism for the acquisition of this cluster at same point in the evolutionary history of these strains. Orthologues of both MULL and MULS, with some rearrangements, were identified in *Subdoligranulum variabile* CIM:MAG 1127, suggesting that mannan utilization could be a metabolic feature shared with other Ruminococcaceae members.

To further understand the distribution of the two MULs within human-associated *F. prausnitzii* strains, we surveyed the publicly available metagenome data from a total of 2441 individuals from regions with distinct geography (North America, Europe, China and Japan) and dietary patterns (Fig. 1b). Overall, 26.35% of the subjects harbor the two *Fp*MULs identified in this study while 37.7% carries the *Fp*MULL, irrespective of the nationality or health state. When examined for frequency within single datasets, different cohorts and nationalities exhibited differing trends. The two *Fp*MULs were most common in the European (up to 40.75% of the subjects), Chinese (up to 48.27%) and Japanese metagenomes (20.37%), whereas their prevalence was lower in North American (10.78%) metagenomes. Among the two hunter-gatherer populations, the Yanomami and Hadza, we detected the presence of only *Fp*MULL in one Yanomami and two Hadza individuals, indicating that these microbiomes may be able to degrade galactomannan derived from tubers that are part of their diet (25).

### *F. prausnitzii* grows efficiently on β-mannooligosaccharides

Growth studies showed that *F. prausnitzii* SL3/3 failed to grow on KGM and CGM (Fig. 2a), likely reflecting the absence of a surface β-1,4-endomannanase required to generate suitable β-mannooligosaccharides (β-MOS) for import into the cell. This hypothesis was confirmed by growing *F. prausnitzii* on both substrates pre-digested with a GH26 β-1,4-endomannanase from *R. intestinalis* (Fig 2a). To assay for oligosaccharide generation and/or uptake, we used HPAEC-PAD and determined the concentration of β-MOS in the initial and spent supernatant from *F. prausnitzii* cultures (Fig 2b). Only polymeric β-mannan was observed in the spent supernatant after growth of the bacterium on KGM and CGM, demonstrating that *F. prausnitzii* does not display surface β-1,4-endomannanase activity. In contrast, *F. prausnitzii* was able to take up and utilize CGM- and KGM-derived β-MOS while mannose and mannobiose (M2) are seemingly untouched.

**Figure 2.**
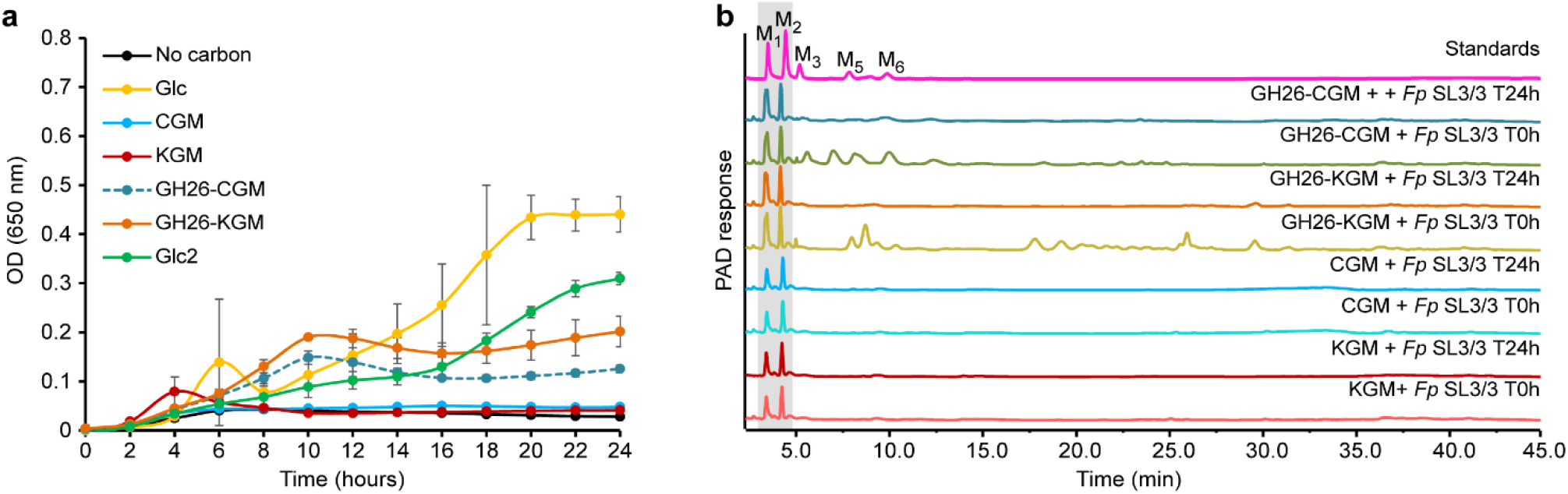
Growth profile and carbohydrate consumption of *F. prausnitzii* SL3/3. **a**, Cells were grown on M2 medium supplemented with 0.2% (w/v) polysaccharide (CGM, carob galactomannan; KGM, konjac glucomannan), oligosaccharides (GH26-CGM, *Ri*GH26-pretreated carob galactomannan; GH26-KGM, *Ri*GH26-pretreated konjac glucomannan), cellobiose (Glc2) and glucose (Glc) as the sole carbon source. Data are averages and standard deviations of three biological replicates. **b**, Analysis of the growth medium used in the experiment described in **a** by HPAEC-PAD. Traces show mannose, mannooligosaccharides and polysaccharides detected in the supernatant before (T0h) and after fermentation (T24h) with *F. prausnitzii*. Samples were chromatographed with the following external standards: M_1_, mannose; M_2_, mannobiose; M_3_, mannotriose; M_4_, mannotetraose; M_5_, mannopentaose; M_6_, mannohexaose. The data displayed are examples from three biological replicates.

Taken together, these data support the concept that the two MULs are being expressed and the resulting proteins orchestrate the degradation of different β-MOS. To determine the biochemical basis for β-MOS import and de-ornamentation, the specificity of β-MOS-binding protein, *Fp*GH36, *Fp*GH113 and the two CEs was determined. A model for catabolism of CGM- and KGM-derived β-MOS is presented in Fig. 3.

**Figure 3.**
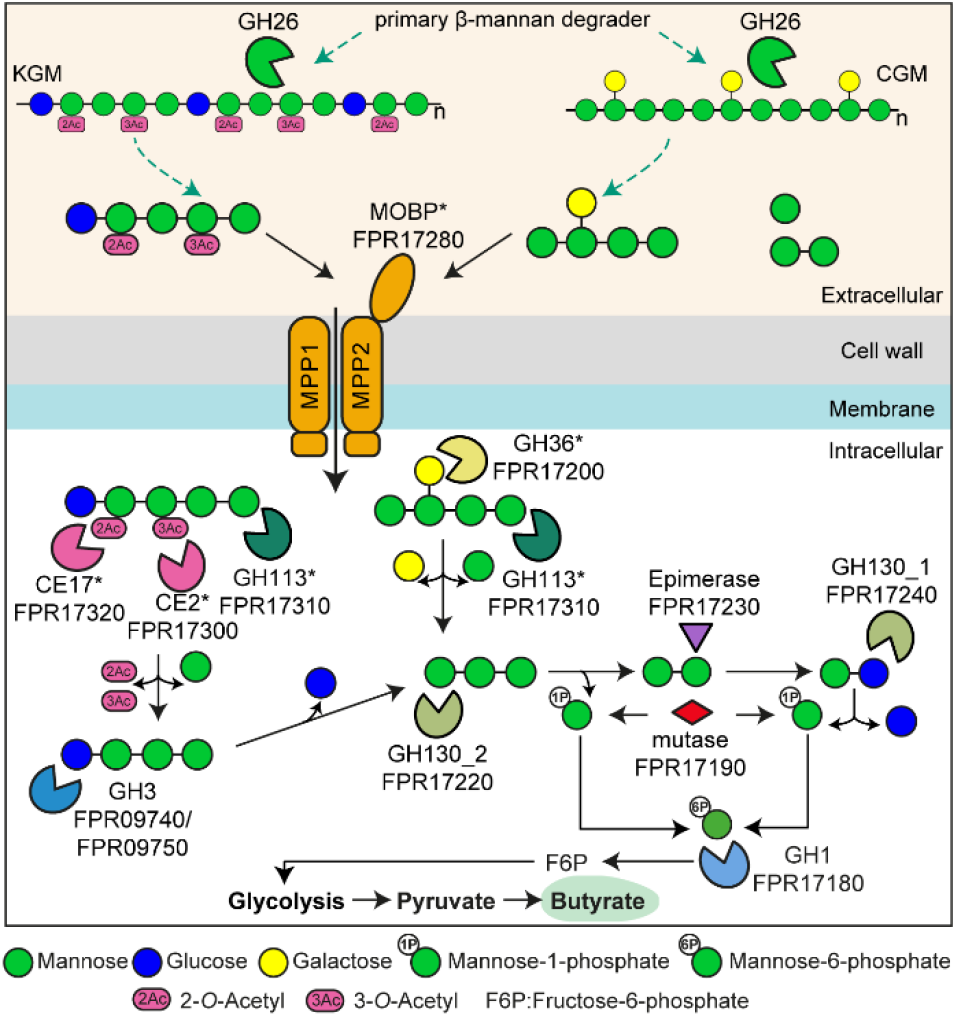
Schematic model of the β-MOS degradation pathway in *F. prausnitzii*. Gene products are colored as in Figure 1. β-MOS liberated by β-mannan keystone species are bound on the surface of the *F. prausnitzii* cells by the MOBP binding protein. The β-MOS transit intracellularly through the associated ABC-transporter. Once intracellularly, the α-galactosidase GH36 and the reducing end mannose-releasing exo-oligomannosidase GH113, respectively, process the galactomannooligosaccharides into galactose, mannose and mannotriose. The glucomannooligosaccharides are processed by the acetylesterases CE2 and CE17, two β-glucosidases (GH3) and the reducing end mannose-releasing exo-oligomannosidase GH113, respectively, into acetate, mannose, glucose and mannotriose. The mannotriose is hydrolyzed by the β-1,4-mannooligosaccharide phosphorylase GH130_2 into mannose-1-phosphate and mannobiose which is then epimerized to mannosyl-glucose by an epimerase. A β-1,4-mannosylglucose phosphorylase GH130_1 phosphorolyses mannosyl-glucose into mannose-1-phosphate and glucose. The mannose-1-phosphate is converted into mannose-6-phosphate by a mannose phosphate mutase and further isomerized into fructose-6-phosphate by a GH1. This product, together with the other liberated monosaccharides and acetate, enters glycolysis that generates pyruvate, some of which is converted into butyrate. Asterisks indicate proteins characterized in this study.

### *Fp*MOBP is a binding protein specific for β-MOS

The binding of mannohexaose and cellohexaose to the recombinantly produced *Fp*MOBP was tested using isothermal calorimetry (ITC). *Fp*MOBP bound to mannohexaose with a K_d_ of 189 ± 1.4 µM (ΔG= −5.08 ± 0.01 kcal/mol; ΔH= −19.8 ± 0.28 kcal/mol; TΔS= 14.7 ± 0.14 kcal/mol; n= 0.8; corresponding thermograms are shown in Fig. S2a). *Fp*MOBP did not show any appreciable binding to cellohexaose (Fig. S2b), demonstrating the specificity of *Fp*MOBP toward mannopyranosyl–linked ligands. Together, these data demonstrate that *Fp*MOBP is part of an ABC transporter specific for β-MOS.

### *Fp*GH113 is a reducing end mannose-releasing exo-oligomannosidase

*Fp*GH113 is a 35 KDa protein sharing 53% identity with *Ri*GH113 from the previously characterized β-mannan utilization system in *R. intestinalis* (7) (Fig. 1). The closest structurally characterized homolog of *Fp*GH113 is the β-1,4-mannanase AxMan113A from *Amphibacillus xylanus* (26) with 48% identities between the two amino acid sequences. No signal peptide was identified by SignalP 4.0, suggesting that *Fp*GH113 is likely located intracellularly. The *Fp*GH113 enzyme released mannose and oligosaccharides from 6^3^,6^4^-α-D-galactosyl-mannopentaose (Gal_2_Man_5_) (Fig. S1a-b) and mannopentaose (Man_5_) (Fig. 4a), with mannose increasing over time (Fig. S1c), consistent with exo-activity. When the reducing end of Man_5_ was reduced with NaBD_4_ (Fig. 4a), no *Fp*GH113 activity could be detected demonstrating that this enzyme is a reducing end mannose-releasing exo-oligomannosidase. Considering the predicted intracellular location of *Fp*GH113, we tested its activity against *Ri*GH26-prehydrolysed CGM. Consistent with this view, release of mannose was detected after overnight incubation of the enzyme with *Ri*GH26-generated galacto-β-MOS (Fig. 4b), while *Fp*GH113 was not able to hydrolyze intact CGM (Fig. S1d).

**Figure 4.**
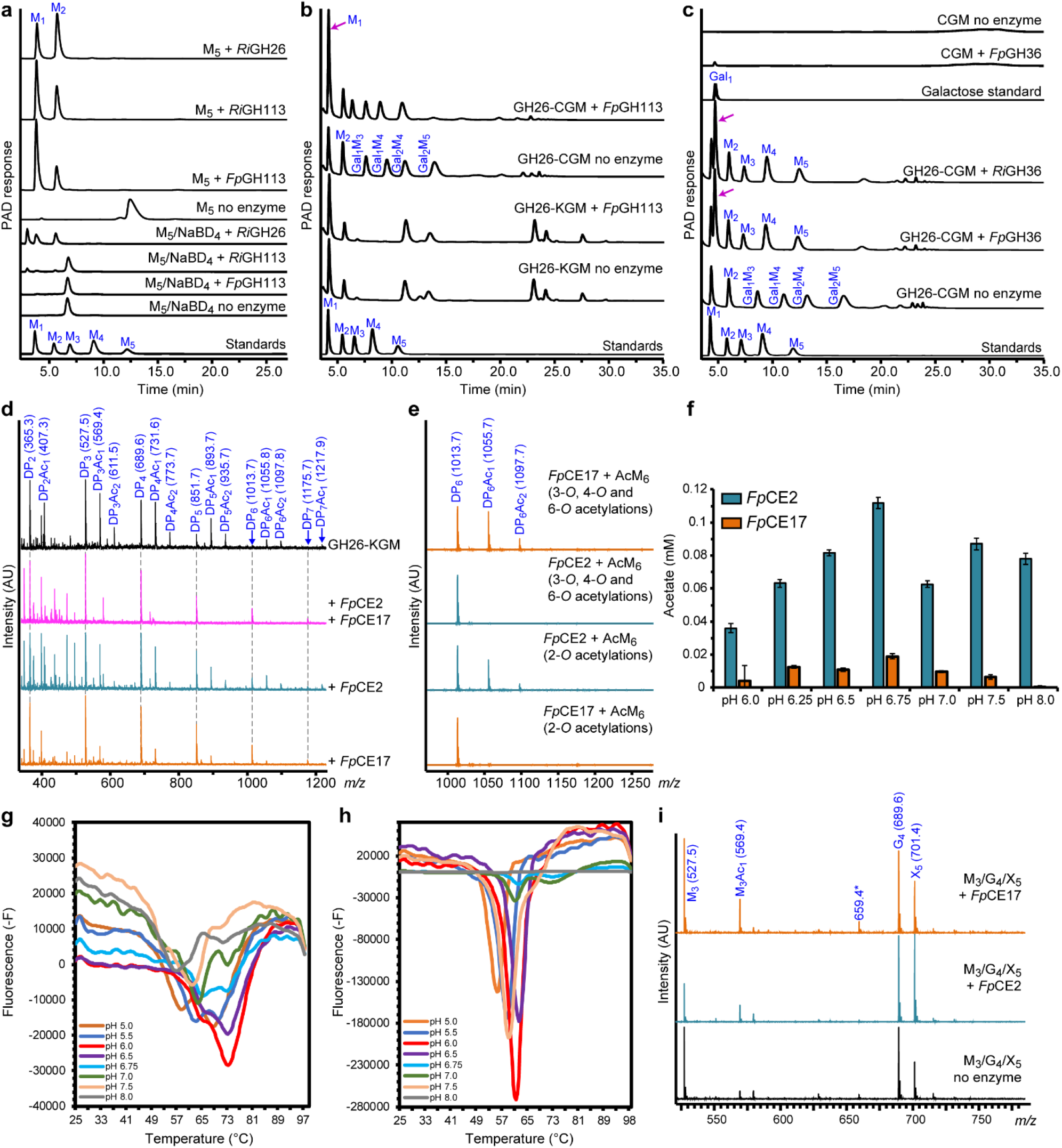
HPAEC-PAD and MALDI-ToF analysis of the activity of enzymes removing acetyl and galactosyl side chains and further hydrolyzing the imported β-MOS. **a**, mannose and β-MOS generated when *Fp*GH113 was incubated with mannopentaose (M_5_). *Fp*GH113 was unable to hydrolyze M_5_ that had been pre-treated with sodium borodeuteride (NaBD_4_), to convert the reducing end monosaccharide unit into its alditol. Control reaction with the previously characterized reducing end mannose-releasing exo-oligomannosidase *Ri*GH113 and endo-mannanase *Ri*GH26 are shown. **b**, HPAEC-PAD traces of the product generated before and after hydrolysis of *Ri*GH26-pretreated carob galactomannan (GH26-carob) and *Ri*GH26-pretreated konjac glucomannan (GH26-KGM) with *Fp*GH113. **c**, HPAEC-PAD trace showing the oligosaccharide products of CGM digestion with *Ri*GH26 and subsequently incubated with *Fp*GH36 α-galactosidase. **d**, MALDI-ToF spectra showing products (as sodium-adducts) generated after incubation of predigested KGM (*Ri*GH26-KGM) with either *Fp*CE17 or *Fp*CE2 or both enzymes in combination. Peaks are labelled by DP and number of acetyl (Ac) groups. **e**, Mass spectra of mannohexaose containing acetylations at different positions after treatment with either *Fp*CE17 or *Fp*CE2. These substrates were generated in house using the *R. intestinalis* esterases *Ri*CE2 and *Ri*CE17. The annotated *m/z* values indicate sodium adducts. **f**, pH optima of *Fp*CE17 or *Fp*CE2. pH optima were determined on pNP-acetate in 50 mM sodium phosphate buffers with varying pH at room temperature. Acetate release was measured after a 10 min incubation. **g**, Thermal shift assay melting curve for *Fp*CE17. **h**, Thermal shift assay melting curve for *Fp*CE2. Both in **g** and **h**, plots show derivative fluorescence data (-F) as a function of temperature (°C). **i**, MALDI-ToF MS analysis of reactions for identification of preferred oligosaccharides for *Fp*CE2 and *Fp*CE17. The esterases were tested on a mix with mannotriose (*m/z* 527; M_3_), cellotetraose (*m/z* 689; G_4_) and xylopentaose (*m/z* 701; X5) and with vinyl acetate, vinyl propionate and vinyl butyrate as ester donors. In all panels, data are representative of independent triplicates. Abbreviations: M_1_, mannose, M_2_, mannobiose; M_3_, mannotriose; M_4_, mannotetraose; M_5_, mannopentaose; M_6_, mannohexaose, Gal_1_, galactose; Gal_1_M_3_, galactosylmannotriose; Gal_1_M_4_, galactosylmannotetraose; Gal_2_M_4_, digalactosylmannotetraose; Gal_2_M_5_, digalactosylmannopentaose.

### Removal of α-galactosyl and acetyl substitutions from β-MOS

*Fp*GH36 is a predicted intracellularly localized 79 KDa enzyme with two GH36 domains, located at the N- and C-terminus of the protein, as well as an internal melibiase domain. The *F. prausnitzii* GH36 domains were all similar to those found in well-characterized α-galactosidases, with AgaB from the thermophilic bacterium *Geobacillus stearothermophilus* being the closest structurally characterized homolog (44 % identity) (27). *Fp*GH36 showed 42% identity to *Ri*GH36 from *R. intestinalis* (Fig. 1). *Fp*GH36 hydrolyzed α-1,6-galactose side-chains from CGM-derived β-MOS (Fig. 4c) and Gal_2_Man_5_ (Fig S1a-b), exhibiting minor activity against polymeric galactomannan (Fig. 4c). This is consistent with the sequential activity of *Fp*GH36 on internalized galacto-β-MOS *in vivo*.

We have previously shown that in *R. intestinalis* the complete removal of acetyl substitutions on the β-MOS backbone is achieved through the complementary action of two esterases, where *Ri*CE2 attacks acetyl groups on either the 3-*O*, 4-*O* or 6-*O* position, while *Ri*CE17 on the 2-*O* position (28). To explore whether *F. prausnitzii* employs a similar mechanism, KGM was pre-hydrolyzed with *Ri*GH26 to generate glucomanno-oligosaccharides (GMOS) that were subsequently incubated with *Fp*CE2 (32% amino acid sequence identity with *Ri*CE2) and *Fp*CE17 (46% amino acid sequence identity with *Ri*CE17). MALDI-ToF MS analysis of products released from GMOS revealed that the two enzymes mediated the complete removal of acetylations when added together while a partial deacetylation was observed when the substrate was treated with each of the enzymes separately (Fig. 4d). To explore the extent to which this strategy for complete substrate deacetylation is conserved in Firmicutes, we exploited the transacetylation specificity of *R. intestinalis* esterases (28) to generate acetylated mannohexaoses (AcM_6_) and tested the activity of *Fp*CE2 and *Fp*CE17 on these substrates. *Fp*CE2 was only able to deacetylate the *Ri*CE2-generated AcM_6_, thus demonstrating that this enzyme removes 3-*O*-, 4-*O*- and 6-*O*-acetylations (Fig. 4e). *Fp*CE17 was effective on *Ri*CE17-generated AcM_6_ and displayed no activity on *Ri*CE2-generated AcM_6_, thus showing that *Fp*CE17 exclusively removes the axially oriented 2-*O*-acetylations (Fig. 4e). Taken together, these results prove that the *F. prausnitzii* esterases have the same acetylation site specificity as their corresponding enzymes in *R. intestinalis*.

To further characterize the two *F. prausnitzii* esterases, we evaluated their activity both on a commercial substrate, i.e. para-Nitrophenyl (pNP) acetate, and on a natural substrate, i.e. *Ri*GH26 hydrolyzed AcGGM. When tested on pNP acetate, both *Fp*CE2 and *Fp*CE17 were most active at pH 6.75 (Fig. 4f). Deacetylation rate measurements on *Ri*GH26-prehydrolyzed AcGGM at pH 6.75 and 35 °C, conditions that prevents acetyl migration, at equal enzyme loadings (50 nM) indicated that *Fp*CE2 releases acetate approximately four times faster than *Fp*CE17 (Table 1). When combined, using 25 nM of each esterase, the deacetylation rate, k_cat_ and specific activity were approximately 2-fold higher compared to the values from treatments with *Fp*CE17 and 2-fold lower compared to the values from treatments with the *Fp*CE2 when used on its own, respectively (Table 1).The reduced resulting rate of deacetylation suggests that the esterases are not acting synergistically but may rather be competing for the substrate, a behavior previously reported in cocktails of multiple enzymes for lignocellulose hydrolysis (29).

**Table 1.** Deacetylation rate, specific activity and turnover rates of *Fp*CE2 and *Fp*CE17 on AcGGM from Norway spruce. Values are calculated based on the acetate released in the initial 15 minutes of reaction.

|  | *FpCE17* | *FpCE2* | *FpCE17+FpCE2* |
| --- | --- | --- | --- |
| **Deacetylation rate [nmol/s]** | 4166 | 16550 | 7588 |
| **$k_{\text{cat}}$ [$\text{s}^{-1}$]** | 83 | 331 | 152 |
| **Specific activity [nmol acetate/s/$\mu\text{g}$ enzyme]** | 2 | 8 | - |

Melting curves for both enzymes in buffers at pH 5.0-8.0 were obtained using a protein thermal shift assay (Fig. 4g-h). Both *Fp*CE2 and *Fp*CE17 displayed an irreversible thermal unfolding transition, which is consistent with their multi-domain structure (28, 30). *Fp*CE17 was stable up to 73 °C, with the highest observed melting temperature at pH 6.0; its lowest observed melting temperature was 58 °C at pH 5. For *Fp*CE2 the unfolding took place at higher temperature, with a melting point of 62 °C at pH 6.0 and a highest melting point of 73 °C at pH 7.0 and 8.0 (Fig. 4g-h); its lowest observed melting temperature was 56 °C at pH 5.0.

Studies of substrate specificities have shown that acetyl esterases are able to efficiently catalyze the transfer of an acetyl group from a donor, such as vinyl acetate, to precise positions of an oligosaccharide with the generation of highly specific esterified oligosaccharides (28). Consistent with that notion, we found that both *F. prausnitzii* esterases were able to transacetylate mannotriose and mannotetraose (data not shown). To further test the preferred substrate for the esterases, we incubated either *Fp*CE17 or *Fp*CE2 with a mix of M_3_, G_4_ and X_5_ and used vinyl acetate as acetyl donor. MALDI-ToF MS analysis of products generated by these reactions showed that the esterases transferred the acetyl group only to M_3_ (Fig 4i), thus confirming the manno-oligosaccharide specificity of *Fp*CE17 and *Fp*CE2.

### Co-cultivation of *F. prausnitzii* with primary β-mannan degraders

The data presented above suggest that *F. prausnitzii* has a sufficiently complex enzymatic toolbox to benefit from the uptake of β-MOS liberated in the surrounding environment by other gut microbes. To test this hypothesis and evaluate the competitiveness of this strain in the utilization of β-MOS, we co-cultured *F. prausnitzii* with two keystone commensal organisms for β-mannan utilization, namely the Gram-negative Bacteroidetes *B. ovatus* strain V975 and the Gram-positive Firmicutes *R. intestinalis* strain L1-82. *F. prausnitzii* grew in the presence, but showed poor growth in the absence, of *B. ovatus* in intact KGM (Fig. 5a). The optical densities obtained when *F. prausnitzii* was grown in monoculture in the no-carbon source control (Fig. 5b) were similar to those obtained in 0.2% (w/v) KGM, suggesting that the microbe is not able to utilize this glycan on its own. Notably, the maximum OD_650_ of the co-culture (OD_650_ = 0.45) appeared higher in the β-mannan polymer than those observed by *B. ovatus* in single culture (OD_650_ = 0.37), indicating that syntrophic growth exists between these two populations in these conditions (Fig. 5a). *F. prausnitzii* is a butyrate producer while carbohydrate fermentation by *B. ovatus* results in the production of propionate (5). Therefore, comparing differences in butyrate levels between the single *F. prausnitzii* culture and co-culture may provide evidence as to whether cross-feeding of β-mannan breakdown products by *F. prausnitzii* occurred. Butyrate concentrations were significantly increased (*p* = 0.004) in the co-culture compared to the mono-culture in KGM (Fig. 5d) or the co-culture in minimal medium (Fig. 5e), which suggests that *F. prausnitzii* can effectively compete for β-MOS generated by the cell-surface exposed endo-mannanase *Bo*Man26B from *B. ovatus* (31). This effect required the presence of living *B. ovatus* cells, as no evidence of an increase of butyrate levels was detected when *F. prausnitzii* was co-grown with a heat-treated *B. ovatus* culture (Fig. S3). When *F. prausnitzii* was co-cultured with *R. intestinalis* in 0.2% KGM or in the absence of a carbon source, the growth curves appeared very similar to when *R. intestinalis* was cultured on its own (Fig. 5g-h). As both bacteria produce butyrate, we compared the value observed in the co-culture to the sum of butyrate concentration in both single cultures. No significant increase (*p* > 0.05) of butyrate concentrations was observed in the co-culture compared to the single cultures in KGM (Fig. 5j) or minimal medium (Fig. 5k). Co-cultivation of *F. prausnitzii* with either *B. ovatus* (Fig. 5c) or *R. intestinalis* (Fig. 5i) on glucose resulted in no increase in the overall levels of butyrate in the co-cultures (Fig. 5f and Fig. 5l). Together with the results observed in the co-cultures grown without any carbon source, these data indicate the specific effect of KGM degradation products to support *F. prausnitzii* growth and exclude the possibility that this microbe is cross-feeding on bacterial derived components (such as capsular polysaccharides).

**Figure 5.**
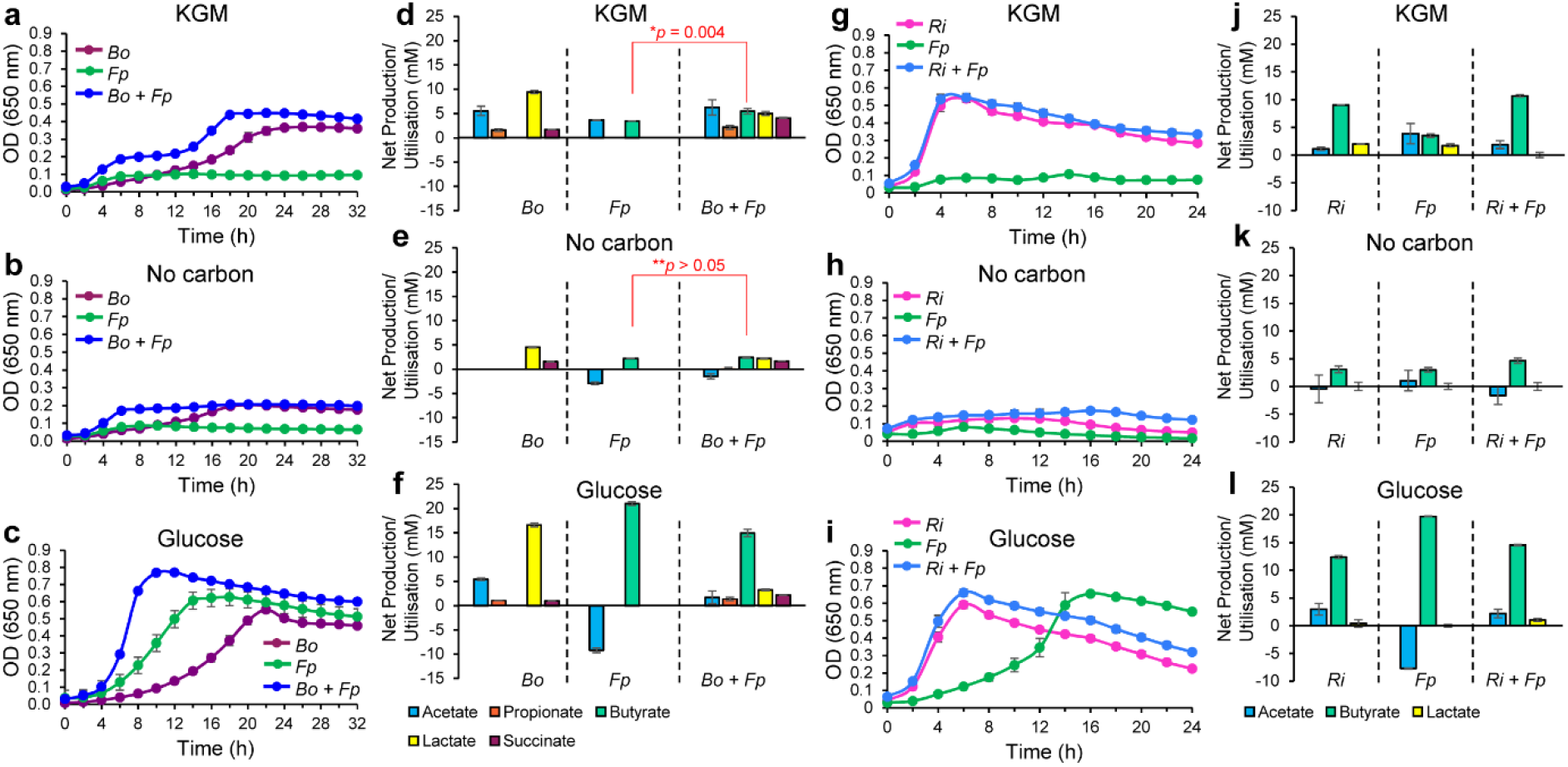
Co-cultivation experiments of *F. prausnitzii* with keystone β-mannan degraders. Growth kinetics of mono- and co-cultures of *F. prausnitzii* (*Fp*) and *B. ovatus* (*Bo*) in M2 medium containing **a**, 0.2% konjac glucomannan (KGM), **b**, no carbon source (no carbon) or **c**, 0.2% glucose. Fermentation products for mono- and co-cultures of *F. prausnitzii* (*Fp*) and *B. ovatus* (*Bo*) in **d**, KGM, **e**, no carbon source or **f**, glucose. Growth of single and co-cultures of *F. prausnitzii* (*Fp*) and *R. intestinalis* (*Ri*) in M2 medium supplemented with **g**, 0.2% konjac glucomannan (KGM), **h**, no carbon source or **i**, glucose. Concentration of different metabolites in the spent media of single and co-cultures of *F. prausnitzii* (*Fp*) and *R. intestinalis* (*Ri*) in M2 medium with **j**, 0.2% konjac glucomannan (KGM) **k**, no carbon source or **l**, glucose. The data are means with standard deviations of biological triplicates. Statistically significant differences were determined using the two-tailed unpaired Student’s t test.

## DISCUSSION

Biochemical work presented herein demonstrates that two MULs support the ability of *F. prausnitzii* to utilize β-MOS. β-MOS from diet are highly variable with respect to sugar composition and linkages and the *F. prausnitzii* enzymatic apparatus is adapted to deal with this diversity. Our findings show that β-MOS are bound by *Fp*MOBP at the cell surface and subsequently imported intracellularly; here, they are further saccharified by *Fp*GH113 and de-galactosylated and de-acetylated by the combined action of *Fp*GH36, *Fp*CE2 and *Fp*CE17 (Fig. 3). By analogy with the model described in *R. intestinalis* (7), putative β-glucosidases of GH3 may confer the removal of terminal glucose residues in gluco-β-MOS prior to depolymerization of the remaining linear β-MOS by the activity of a putative mannooligosaccharide phosphorylase (*Fp*GH130_2) into mannobiose. Mannobiose is subsequently epimerized into mannosyl-glucose by a putative epimerase, *Fp*MEP and phosphorolysed by *Fp*GH130_1 into glucose and mannose-1-phosphate, similar to the pathway described in *Ruminococcus albus* (32). A comparative genomic analysis revealed that these MULs are widespread and highly conserved amongst human gut-associated *F. prausnitzii* (Fig. 1A). Presence of genes associated with conjugation and phage-related events in the flanking regions suggests that the *Fp*MULL was acquired through horizontal gene transfer from other gut bacteria, as previously observed for PULs identified in commensal *Bacteroides* genomes (33).

Members of the dominant *Bacteroides* genus, such as *B. ovatus*, and *Roseburia* species that possess GH26 endo-mannanases have been described as the keystone bacteria for mannan degradation in the gut (7, 21, 31). In contrast, *F. prausnitzii* may only access oligosaccharides, released by these primary degraders, which can be imported without the need for extracellular enzymatic cleavage. In this context, we demonstrate that β-MOS are indeed released into the culture medium by *B. ovatus* during co-growth on KGM, and that *F. prausnitzii* is capable to efficiently compete for and utilize these oligosaccharides (Fig. 5a and 5d). However, we observed no explicit evidence of cooperative growth between *R. intestinalis* and *F. prausnitzii* (Fig. 5g and 5j). We recently demonstrated the competitiveness of *R. intestinalis* on β-mannan when in co-culture with *B. ovatus* during growth on AcGGM (7), and highlighted a pivotal role of a transport protein (*Ri*MnBP) within the uptake system, which exhibited strong binding to short β-MOS (DP 3-6) with different sidechain decoration patterns (7). Similar observations were recently reported in the *Bifidobacterium* genus, with *B. animalis* subsp. *lactis* ATCC 27673 outcompeting *B. ovatus* during growth on galactomannan (34). Notably, the two *B. animalis* binding proteins, BlMnBP2 and BlMnBP1, mediated high affinity capture of β-MOS with preference to oligosaccharides of DP 3-4 and K_d_ values in the 70-80 nM range (34). In contrast, the SusD-like β-MOS binding proteins from *B. ovatus* displayed binding to M6 (K_d_ value of 1.8 ± 0.2 mM) (31) with about 10-fold lower affinity than that of *Fp*MOBP, 53-fold lower affinity than that of *Ri*MnBP, and about 900-fold lower affinity than that of BlMnBP2 and BlMnBP1. Thus, the differential transporter’ affinity to β-MOS provides a possible rationale for the trophic interactions established by *F. prausnitzii* with *Bacteroides*, enabling efficient capture of communally available nutrients within synthetic consortia, and potentially in natural gut communities. However, the binding affinity of MnBPs plausibly gives reason to why cross-feeding on KGM was unlikely to exist between *R. intestinalis* and *F. prausnitzii*, as the affinity of *Ri*MnBP to M6 (K_d_ value of 33.75 ± 0.95 µM) was five times stronger than that of *Fp*MOBP. This difference is likely to be crucial for selfish resource capture by the keystone β-mannan primary degrader *R. intestinalis* (7), with minimal loss in a competitive environment.

To understand the capacity of human populations to derive nutrition from β-MOS, we surveyed 2441 publicly available human metagenomes and revealed that MULs closely related to those of *F. prausnitzii* are widely distributed throughout human populations (Fig. 1c). Indeed, we did not observe correlation with any particular population or nation, consistent with the fact that dietary β-MOS/β-mannan are a ubiquitous component of the human diet. The *Fp*MULs were more common than population restricted traits like red algal porphyran degradation, known to be confined to a small cohort of Japanese subjects and absent in the microbiome of western individuals (33), but they were less common than *Bacteroides*-associated PULs for degradation of plant cell-wall xyloglucan (92% of samples) (35), mixed-linkage β-glucans (92.5% of samples) (36), β-(1,3)-glucans (59% of samples) (37) and yeast α-mannans (62% of samples) (38). Moving beyond the human microbiota, we detected two analogous *Fp*MULs in a *F. prausnitzii* strain found in the porcine gut microbiota (39). Proteomic analysis identified *Fp*MUL-encoded proteins being more abundant in pigs fed a diet supplemented with 4% acetylated galactoglucomannan, thus providing evidence that these analogous MULs are employed by *F. prausnitzii* inhabiting environments beyond the human gut (39).

In conclusion, biochemical and microbiological data presented in this study illustrate that *F. prausnitzii* possesses an extensive enzymatic apparatus that targets β-MOS released by neighboring colonic bacteria. ITC data provide evidence that the external recognition machinery is tailored for the capture of β-MOS with stronger affinity than *Bacteroides*. This is in line with the fact that, when in co-culture, *F. prausnitzii* showed cross-feeding behaviors with *B. ovatus*, whose own β-MOS uptake requires a SusD-like protein that binds oligosaccharides with about 10-fold lower affinity than *Fp*MOBP. Furthermore, this study in conjunction with a previous report (7) points to a competitive mechanism of β-MOS/β-mannan utilization in the gut microbiota where keystone Lachnospiraceae members like *R. intestinalis* have developed a sophisticated “selfish” uptake and degrading system to minimize sharing resources with *Bacteroides* and other Ruminococcaceae species such as *F. prausnitzii*.

Overall, our study contributes towards the understanding of cross-feeding mechanisms deployed by a beneficial commensal organism to interact with dietary β-mannan. Significantly, these findings could help to design prebiotic/symbiotic formulations that are optimized for selective manipulation of gut microbiome functions in ways that promote human health and beyond.

## MATERIALS AND METHODS

### Substrates

All glycan stocks were prepared at 10 mg/ml in ddH_2_O and sterilized by filtration using a 0.22 µm membrane filter (Sarstedt AG & Co, Germany).

#### Polysaccharides

Konjac glucomannan and carob galactomannan were purchased from Megazyme International (Wicklow, Ireland).

#### Oligo- and monosaccharides

Mannose (M_1_) and glucose (G_1_) were purchased from Sigma Aldrich (St. Louis, MO, USA). Xylopentaose (X_5_), cellobiose (G_2_), cellotetraose (G_4_), mannobiose (M_2_), mannotriose (M_3_), mannotetraose (M_4_), mannopentaose (M_5_), mannohexaose (M_6_), 6^1^-α-D-galactosyl-mannotriose (GalM_3_) and 6^3^, 6^4^-α-D-galactosyl-mannopentaose (Gal_2_M_5_) were purchased from Megazyme. Konjac glucomannan digest and carob galactomannan digest were produced in-house using *Ri*GH26 (7) in 10 mM sodium phosphate, pH 5.8. Reactions were incubated for 16 h at 37 °C following removal of *Ri*GH26 using a Vivaspin 20 filtration unit (10.000 MWCO PES, Sartorius) and carbohydrate lyophilization on an ALPHA 2-4 LD Plus freeze dryer (Christ, Germany). Acetylated galactoglucomannan (AcGGM) was produced in-house as described by La Rosa et al. (24).

### Bacterial strains and culture conditions

*F. prausnitzii* SL3/3, *R. intestinalis* L1-82 and *Bacteroides ovatus* V975 were routinely cultured under CO_2_ at 37 °C in M2 medium containing 30% clarified rumen fluid supplemented with 0.2% (w/v) glucose, soluble potato starch and cellobiose (GSC) (40). Growth measurements on individual substrates were performed in M2GSC medium containing a single carbohydrate at 0.2% (v/v) final concentration using 96-well plates in a Don Whitley MACS-VA500 workstation (80% N_2_, 10% H_2_, 10% CO_2_). Growth was assessed by measuring the absorbance at 650 nm (OD_650_) at 2 h intervals for up to 24 h using an Epoch 2 microplate reader (BioTek, Vermont, USA). The competition assay of *F. prausnitzii* and either *B. ovatus* or *R. intestinalis* were conducted by growing the strains as above in the presence of 0.2% (v/v) konjac glucomannan. The strains inoculated into M2 medium with no added carbohydrate source and M2 medium with 0.2% (v/v) glucose were included as negative and positive controls, respectively. Five µl of overnight bacterial cultures from both strains were used to inoculate the wells (final volume of 200 µl). The co-cultures were incubated at 37 °C anaerobically and growth was followed by measuring the OD_650_ for 24-32 h. Samples (500 µl) were collected at the end of the experiment for SCFAs analysis. All growth experiments were performed in triplicates.

### Cloning, expression, and purification of recombinant protein

The genes encoding mature forms of the proteins described in this study were amplified from the *F. prausnitzii* SL3/3 genomic DNA (BioProject accession number PRJNA39151) by PCR, using appropriate primers (Table S1). All primers were designed to amplify constructs to exclude predicted signal peptides (predicted by the SignalP v4.1 server (41)). PCR products were generated using the Q5 High-Fidelity DNA Polymerase (New England BioLabs, United Kingdom) with 50 ng genomic DNA as template. The PCR products were cloned into pNIC-CH (Addgene plasmid 26117) by ligation-independent cloning (42). All constructs were designed to harbor a C-terminal His_6_-tag fusion in the translated recombinant peptide although, for *Fp*GH36, His-tag translation was prevented by the introduction of one stop codon at the end of the open-reading frame. Successful generation of constructs was verified by sequencing (Eurofins, UK). Plasmids harboring the gene of interest were transformed into chemically competent *E. coli* BL21 STAR cells (Invitrogen) and an overnight pre-culture was inoculated to 1% in 500 ml Tryptone Yeast extract (TYG) containing 50 μg/ml kanamycin, followed by incubation of the fresh culture for 16 h at 25 °C. Protein overexpression was induced by adding isopropyl β-D-thiogalactopyranoside (IPTG) to a final concentration of 200 μM. Recombinant protein production continued overnight at 25 °C, after which the cells were collected by centrifugation. *Fp*CE17, *Fp*CE2, *Fp*GH113 and *Fp*MOBP were purified by immobilized metal ion affinity chromatography (IMAC). To this aim, the harvested cell pellet was resuspended in binding buffer (20 mM sodium phosphate pH 7.4, 500 mM sodium chloride, 5 mM imidazole) and lysed using a Vibracell Ultrasonic Homogenizer (Sonics and Materials, USA). The cell debris was pelleted by centrifugation and the supernatant was loaded onto a 5 ml HisTrap IMAC HP Nickel Sepharose column (GE Healthcare), using an ÄKTA Pure chromatography system (GE Healthcare). The target His-tagged protein was eluted using a linear gradient of 0–100% elution buffer (20 mM sodium phosphate pH 7.4, 500 mM sodium chloride, 500 mM imidazole) over 16 column volumes. *Fp*GH36 was purified by hydrophobic interaction chromatography (HIC). *Fp*GH36-containing cell pellet was resuspended in a buffer with 1.5 M ammonium sulfate and lysed as described above. The cell free supernatant was loaded onto a 5 ml HiTrap Phenyl FF (GE Healthcare) and protein was eluted by using a linear reverse gradient to 100 mM NaCl over 90 min at a flow rate of 2.5 ml/min. After IMAC or HIC purification, eluted protein fractions were pooled, concentrated using a Vivaspin 20 centrifugal concentrator (10-kDa molecular weight cutoff) and applied to a HiLoad 16/600 Superdex 75 pg gel filtration column (GE Healthcare). Pure protein samples were dialyzed against 10 mM TrisHCl pH 7.0 and concentrated as described above. Protein purity was determined by SDS-PAGE analysis. Protein concentrations were determined using the Bradford assay (Bio-Rad, Germany).

### Activity assays

Unless otherwise stated, enzyme reactions contained 10 mM sodium phosphate, pH 5.8, and 0.1 mg/ml substrate. Reactions were pre-heated (37 °C for 10 min) in a Thermomixer C incubator with a heated lid (Eppendorf), before addition of the enzyme to 1 μM (in a final reaction volume of 100 μl) for further incubation (up to 16 h) at 37 °C and 700 rpm. All experiments were performed in triplicates.

### MALDI-ToF MS analysis of oligosaccharides

Mannooligosaccharides products were analyzed by MALDI-ToF mass spectrometry on a Ultraflex MALDI-ToF/ToF MS instrument (Bruker Daltonics, Germany) equipped with a 337-nm-wavelength nitrogen laser and operated by the MALDI FlexControl software (Bruker Daltonics). A matrix of 2,5-dihydroxybenzoic acid (DHB) (9% 2,5-dihydroxybenzoic acid [DHB] – 30% acetonitrile [v/v]), was used. All measurements were performed in positive ion, reflector mode with 1000 shots taken per spectrum.

### Carbohydrate analysis using high-performance anion-exchange chromatography

Oligo- and monosaccharides were analyzed by high-performance anion-exchange chromatography with pulsed amperometric detection (HPAEC-PAD) on a Dionex ICS-3000 system operated by Chromeleon software version 7 (Dionex). Sugars were loaded onto a CarboPac PA1 2 × 250-mm analytical column (Dionex, Thermo Scientific) coupled to a CarboPac PA1 2 × 50 - mm guard column kept at 30 °C. Depending on the analytes, the following gradients were used. The system was run at a flow rate of 0.25 ml/min. For manno-oligosaccharides, the elution conditions were 0-9 min 0.1 M NaOH; 9-35 min 0.1 M NaOH with a 0.1 to 0.3 M NaOAc gradient; 35-40 min 0.1 M NaOH with 0.3 M NaOAc; and 40-50 min 0.1 M NaOH. Commercial mannose and mannno-oligosaccharides (DP 2 to 6) were used as external standards.

### Acetate release measurements using high-performance liquid chromatography

Acetate content in the samples was analyzed on an RSLC Ultimate 3000 (Dionex, USA) HPLC using a REZEX ROA-Organic Acid H+ 300 × 7.8 mm ion exclusion column (Phenomenex, USA). The injection volume was 5 μl and separation was conducted at 65°C, with isocratic elution using 0.6 ml/min of 5 mM H_2_SO_4_ as mobile phase. The UV detector was set to 210 nm. Data collection and analysis were carried out with the Chromeleon 7.0 software (Dionex).

### SCFAs analysis

SCFA concentrations were measured using a gas chromatograph analyzer equipped with a flame ionization detector (GC-FID) as described previously (43). Following derivatization of the samples using N-tertbutyldimethylsilyl- N-methyltrifluoroacetamide, the samples were analyzed on a Hewlett-Packard 6890 gas chromatograph equipped with a silica capillary column using helium as the carrier gas. Quantification of SCFA in the chromatograms was determined based on the retention times of the respective SCFA standards (Sigma-Aldrich, United Kingdom) at concentrations ranging between 5 to 30 mM.

### Isothermal titration calorimetry

Binding of mannohexaose and cellohexaose to *Fp*MOBP was measured at 25 °C in 50 mM sodium phosphate pH 6.5 using either a MicroCal ITC_200_ microcalorimeter or a MicroCal VP-ITC system. To assess the binding to mannohexaose using a MicroCal ITC_200_ microcalorimeter, *Fp*MOBP in the sample cell (2.5 µM) was titrated by a first injection of 0.5 μl followed by 19 × 2 μl injections of carbohydrate ligand (2.5 mM) with 120 s between injections. To evaluate the binding to cellohexaose using a MicroCal VP-ITC system, *Fp*MOBP in the sample cell (22.5 µM) was titrated by a first injection of 2 μl followed by 29 × 6 μl injections of carbohydrate ligand (2.5 mM) with 180 s between injections. Thermodynamic binding parameters were determined using either the MicroCal Origin software (version 7.0) or the VPviewer2000 software (version 2.6).

### *Fp*CE2 and *Fp*CE17 optimal pH

The pH optima for *Fp*CE17 and *Fp*CE2 were assessed by incubation of the enzymes with 0.5 mM 4-nitrophenyl (pNP) acetate (Sigma-Aldrich, Germany) at 25 °C using 50 mM sodium phosphate buffer at pHs ranging from 5.0 to 8.0. Due to the difference in deacetylation rate of pNP acetate by the two enzymes, 1 nM of *Fp*CE2 and 0.1 μM of *Fp*CE17 were used in these experiments. Standard plots of 4-Nitrophenol (p-Nitrophenol, Sigma-Aldrich) were prepared at each pH. The experiments were conducted in triplicate in a volume of 100 μL of sample mixture in 96-well microtiter plates. The reaction was followed by measuring the absorbance at 405 nm at 1-minute intervals for 10 minutes using a Microplate reader (BioTek, USA).

### Protein thermal shift assay

The thermal stability of *Fp*CE17 and *Fp*CE2 was examined using the Protein Thermal Shift™ Kit (ThermoFisher, USA) by measuring fluorescence in a real-time PCR system (Applied Biosystems™, USA). A final concentration of 0.1 mg/ml of *Fp*CE17 and *Fp*CE2 was used in 50 mM sodium phosphate buffers at pH 5.0 to 8.0 and mixed with ROX dye according to the kit protocol. The melting temperature was executed in four replicates with temperatures from 25 to 99 °C in 1 % increments. The data was processed using the StepOne™ software (Applied Biosystems™, USA).

### Transesterification reactions

Transesterification of oligosaccharides was conducted using vinyl acetate (Thermo scientific, USA) as acetate donors. Enzymes (1 µg/ml final concentration) were mixed with 1 mg/ml oligosaccharides and a volume of vinyl acetate corresponding to 50 % of the sample volume was added. The samples were incubated in a thermomixer (Eppendorf, Norway), shaking at 600 rpm, at ambient temperature overnight, then kept at −20 °C until frozen. The vinyl acetate, which remained in liquid phase on top of the frozen aqueous phase, was discarded; enzyme deactivation and carbohydrate precipitation was achieved by adjusting the aqueous phase to 80% (v/v) ethanol with ice-cold 96% ethanol. Enzymes were removed through filtration using a 1 ml Amicon Ultracel 3kDa ultrafiltration device (Merck KGaA, Germany). The samples were then dried using an Eppendorf Concentrator plus (Eppendorf, Norway) at 30 °C and the material dissolved in 100 μl dH_2_O.

### Comparative genomics analysis

Searches for the presence of MULL and MULS in other publicly available *F. prausnitzii* genomes were conducted using a similar strategy as described previously (7). Briefly, the identification of similar MULs in strains other than *F. prausnitzii* SL3/3 was done using BLASTN and the Gene Ortholog Neighborhood viewer on the Integrated Microbial Genomes website (https://img.jgi.doe.gov) using the sequences of the genes coding for *Fp*MOBP (FPR_17280), *Fp*GH113 (FPR_17310) and *Fp*GH3 (FPR_09740) as the search homolog and the default threshold e-value of 1e-5. If this generated a hit, we repeated the process with the adjacent gene to verify that the locus was found in the identified strain. Then, the amino acid identities between each *F. prausnitzii* SL3/3 MULL-MULS RefSeq annotated protein and the hits identified in other *F. prausnitzii* strains were determined by BLASTP-based analyses. Finally, we compared the genomic regions surrounding each orthologous MUL for gene conservation and amino acid identities.

### Analysis of human gut metagenomic data sets for the presence of MULs

Available cohorts of human gut metagenomic sequence data (National Center for Biotechnology Information projects: PRJNA422434 (44), PRJEB10878 (45), PRJEB12123 (46), PRJEB12124 (47), PRJEB15371 (48), PRJEB6997 (49), PRJDB3601 (50), PRJNA48479 (20), PRJEB4336 (51), PRJEB2054 (52), PRJNA392180 (53), and PRJNA527208 (54)) were searched for the presence of MUL nucleotide sequences from *F. prauznitzii* MULL (17.5 Kb) and *F. prauznitzii* MULS (5.5 Kb) using the following workflow. Each MUL nucleotide sequence was used separately as a template and then Magic-BLAST (55) v1.5.0 was used to recruit raw Illumina reads from the available metagenomic datasets with an identity cutoff of 97%. Next, the alignment files were used to generate a coverage map using bedtools (56) v2.29.0 to calculate the percentage coverage of each sample against each individual reference. We considered a metagenomic data sample to be positive for a particular MUL if it had at least 70% of the corresponding MUL nucleotide sequence covered.

## Data availability

All data supporting the findings of this study are available within the article and Supplementary Information.

## Acknowledgements

We are grateful for support from The Research Council of Norway (FRIPRO program to PBP: 250479; BIONÆR program to BW: 244259), as well as the European Research Commission Starting Grant Fellowship (awarded to PBP; 336355 MicroDE).

## Author contributions

S.L.L.R. generated constructs, performed recombinant protein production and purification and functional characterizations of the binding protein and GHs. L.J.L., S.L. and L.M. expressed, purified and performed functional characterization of *Fp*CE2 and *Fp*CE17. Growth experiments on mannans and SCFA quantifications were performed by G.L. ITC was performed by Å.K.R., Z.L. and L.S.M. G.P. and S.L.L.R. conducted the human metagenomic analysis. S.L.L.R., P.B.P and B.W. conceived the study and supervised research. The manuscript was written primarily by S.L.L.R. with contributions from P.B.P., S.D., G.L, L.M., S.L., G.P., E.M., L.S.M., B.W. and L.J.L. Figures were prepared by S.L.L.R.

